# High sensitivity mapping of cortical dopamine D2 receptor expressing neurons

**DOI:** 10.1101/349100

**Authors:** Jivan Khlghatyan, Clémentine Quintana, Martin Parent, Jean-Martin Beaulieu

**Affiliations:** Department of Pharmacology & Toxicology, University of Toronto, Medical Sciences Building, Toronto, ON M5S 1A8, Canada; Department of Psychiatry and Neuroscience, Faculty of Medicine, Université Laval, Québec-City, QC, Canada G1J 2G3

**Author notes:** Correspondence and request for material should be sent to Jean-Martin Beaulieu, Department of Pharmacology & Toxicology, University of Toronto, Medical Sciences Building, Toronto, ON M5S 1A8, Canada. These authors contributed equally to this work (Jivan Khlghatyan, Clémentine Quintana).

**Keywords:** Dopamine D2 receptor, RiboTag, translatome, binary reporter mouse, orphan receptors, prefrontal cortex, claustrum, insula, sensory cortices.

## Abstract

Cortical D2 dopamine receptor (*Drd2*) have mostly been examined in the context of cognitive function regulation and neurotransmission modulation of medial prefrontal cortex by principal neurons and parvalbumin positive, fast-spiking, interneurons in schizophrenia. Early studies suggested the presence of D2 receptors in several cortical areas, albeit with major technical limitations. We used combinations of transgenic reporter systems, recombinase activated viral vectors, quantitative translatome analysis and high sensitivity *in situ* hybridization to identify D2 receptor expressing cells and establish a map of their respective projections. Our results identified previously uncharacterized clusters of D2 expressing neurons in limbic and sensory regions of the adult mouse brain cortex. Characterization of these clusters by translatome analysis and cell type specific labeling revealed highly heterogeneous expression of D2 receptors in principal neurons and various populations of interneurons across cortical areas. Transcript enrichment analysis also demonstrated variable levels of D2 receptor expression and several orphan G-protein-coupled receptors co-expression in different neuronal clusters, thus suggesting strategies for genetic and therapeutic targeting of D2 expressing neurons in specific cortical areas. These results pave the way for a thorough reexamination of cortical D2 receptor functions, which could provide information about neuronal circuits involved in psychotic and mood disorders.

## Introduction

The dopamine D2 receptor (Drd2) is a G-protein coupled receptor (GPCR) and a direct target of antipsychotics that may also contribute indirectly to the action of mood stabilizers (Seeman P et al. 1975; Creese I et al. 1976; Strange PG 2001; Beaulieu JM and RR Gainetdinov 2011; Li X et al. 2012; Del’ Guidice T and JM Beaulieu 2015; Ashok AH et al. 2017). In line with this, *DRD2* gene variants have been identified as risk factors for schizophrenia (Consortium SWGotPG 2014). The receptor has been shown to signal through Gαi/o to inhibit cAMP production (Kebabian JW and DB Calne 1979; Ohara K et al. 1988) and beta-arrestin-2 to inhibit Akt activity (Beaulieu JM et al. 2005). *Drd2* is expressed under two splice variants, termed D2L and D2S (Guiramand J et al. 1995). The highest expression of D2L occurs in striatum (STR) medium spiny neurons and cholinergic interneurons (Missale C et al. 1998; Gerfen CR 2000), while D2S is expressed mostly in dopamine neurons of the *substantia nigra* (SN) and ventral tegmental area (VTA) (Guiramand J *et al.* 1995).

Cortical functions of Drd2 are of interest, considering the involvement of dopamine transmission in the regulation of behavioral dimensions including fear, anxiety, cognition, attention, working memory and information processing in schizophrenia and mood disorders (Seamans JK and CR Yang 2004; Durstewitz D and JK Seamans 2008; Cervenka S et al. 2012; Brisch R et al. 2014; Tovote P et al. 2015; Wang Y et al. 2016; Plavén-Sigray P et al. 2017). A majority of studies have focused on the role of Drd2 in prefrontal cortex pyramidal neurons and interneurons, particularly the parvalbumin (PV) subtype because of their potential contribution to cognitive dysfunctions in schizophrenia (Seamans JK and CR Yang 2004; Santana N et al. 2009; Gee S et al. 2012; Lewis DA et al. 2012; Tritsch NX and BL Sabatini 2012; Murray AJ et al. 2015; Choi SJ et al. 2017). However, early work using radioactively labeled Drd2 antagonist raclopride has demonstrated the presence of D2 binding regions throughout cortex (Lidow MS et al. 1989). Expression of Drd2 in different cortical areas have also been reported (Vincent SL et al. 1993; Gaspar P et al. 1995; Le Moine C and P Gaspar 1998; Santana N *et al.* 2009; Zhang ZW et al. 2010; Gee S *et al.* 2012; Tritsch NX and BL Sabatini 2012), albeit with severe technical limitations due to low mRNA expression and selectivity of antibodies or reporter systems (Tritsch NX and BL Sabatini 2012). Thus, a systematic multimodal, cortex-wide characterization of Drd2 expression still has to be performed to clarify the nature and abundance of Drd2 expressing cells in different cortical regions.

To overcome previous technical limitations we used mice expressing a Cre activated RiboTag, specifically in Drd2 positive (Drd2+) cells (Sanz E et al. 2009; Puighermanal E et al. 2015). RiboTag mice carry a ribosomal protein Rpl22 allele with a floxed wild-type C-terminal exon followed by an identical C-terminal exon that has three HA epitopes inserted before the stop codon. Notably, Rpl22-HA is expressed under the control of its own promoter. Thus, in contrast to analog systems in which reporter protein expression is proportional to Drd2 promoter activity, these mice allow for a highly sensitive, all or none (binary) detection of Drd2+ neurons throughout the brain. Furthermore, the possibility to perform cell-specific translational profiling (Brichta L et al. 2015) provides a direct quantification of endogenous Drd2 mRNA translation in neurons expressing the RiboTag reporter.

Our results revealed previously uncharacterized clusters of Drd2+ cells in limbic and sensory cortical regions of the adult mouse brain. Further analysis quantitatively and qualitatively validated the presence of Drd2 mRNA as well as indicated molecular and cellular heterogeneity of Drd2+ clusters across the cortex. Projection tracing using virally encoded reporters was also used to establish a general map of Drd2 neurons connections that may provide insights into the roles of these neurons. Overall, these findings represent a useful resource to guide investigations of cortical Drd2 functions, Drd2 and associated receptors targeting therapies, while providing valuable information about Drd2 regulated neuronal circuits that are potentially involved in psychotic and mood disorders.

## Material and Methods

### Animals

D2Cre heterozygous bacterial artificial chromosome (BAC) transgenic mice (Gerfen CR et al. 2013) (GENSAT RRID: MMRRC_017263-UCD) were used for virus injections. Homozygous knockin RiboTag (RiboHA) mice (Sanz E *et al.* 2009) were crossed to D2Cre mouse line to generate RiboHA/D2Cre mice and used for ribosome-associated mRNA isolation and immunohistochemistry. BAC D2EGFP mice (Tg(Drd2-EGFP)S118Gsat, MGI:3843608) were used for immunohistochemistry. Mice were maintained on a 12-hours light/dark cycle with ad libitum access to food and water. All experiments conducted in this study are approved by Université Laval and University of Toronto Institutional Animal Care Committee in line with guidelines from Canadian Council on Animal Care.

### AAV viruses

All AAV5 viral particles (AAV-hSyn-GFP; AAV-hSyn-DIO-mCherry) were purchased from University of North Carolina Vector core facility (Chapel Hill, NC).

### Mouse stereotaxic surgery

Mice were anesthetized with a preparation of ketamine 10mg/ml and xylazine 1mg/ml (0.1ml/10g, i.p.). The animal was placed in a stereotaxic frame, and the skull surface was exposed. Two holes were drilled at injection sites and virus (a mixture of AAV-hSyn-GFP and AAV-hSyn-DIO-mCherry) was injected using an injector with microsyringe pump controller (WPI) at the speed of 4nl per second. Following injection coordinates were used:

|  | mPFC | Ant<br>CL/INS | Med<br>CL/INS | Post<br>CL/INS | SS | AUD | VIS |
| --- | --- | --- | --- | --- | --- | --- | --- |
| anteroposterior | 2.4 | 2.1 | 0.9 | -0.3 | -0.8 | -1.8 | -2.8 |
| mediolateral | ±0.5 | ±2.3 | ±3.5 | ±4.3 | ±3.7 | ±4.3 | ±2.9 |
| dorsoventral | -1.7 | -3 | -2.9 | -2.9 | -0.5 | -0.5 | -0.5 |

All measures were taken before, during, and after surgery to minimize animal pain and discomfort.

### Tissue dissection

Mice were killed by rapid cervical dislocation. Heads of animals were immediately cooled by immersion in liquid nitrogen for 6 seconds. For microarray analyses, mPFC tissues were dissected rapidly (within 30 s) on an ice-cold surface and frozen in liquid nitrogen (5 mice per group). For other experiments, first, 500um thick serial coronal sections were prepared using ice-cold adult mouse brain slicer and matrix (Zivic instruments), second, mPFC, anterior claustrum and insular cortex (CL/INS), medial CL/INS, posterior CL/INS, somatosensory cortex (SS), auditory cortex (AUD) and visual cortex (VIS), Endoperiform cortex, Hippocampus and Striatum were dissected on ice cold surface using microsurgical knife (KF Technology).

### Immunoprecipitation of polyribosomes and RNA isolation

Immunoprecipitation of polyribosomes was performed as described before (Sanz E *et al.* 2009). Tissue samples were lysed in homogenization buffer (50mMTris, pH 7.5, 100 mM KCl, 12 mM MgCl2, 1% Nonidet P-40, 1 mM DTT, 100U/mL RNase Out, 100 μg/mL cycloheximide, Sigma protease inhibitor mixture) followed by centrifugation for 10 min at 10000g. Anti-hemagglutinin (HA) antibody (1:150; MMS-101R; BioLegend) was added into collected supernatant and tubes were kept under constant rotation for 4 hours at 4°C. Protein G magnetic beads (Life Technologies) were washed 3 times with homogenization buffer then added into the mixture and kept for constant rotation overnight at 4°C. The following day magnetic beads were washed three times with high salt buffer (50mMTris, pH 7.5, 300 mM KCl, 12 mM MgCl2, 1% Nonidet P-40, 1 mM DTT, 100U/mL RNase Out, 100 μg/mL cycloheximide, Sigma protease inhibitor mixture). RNA was extracted by adding TRI reagent (Zymo research) to magnetic beads pellet followed by Direct-zol RNA kit according to the manufacturer’s instructions (Zymo research). RNA concentration was quantified using ND-1000 Spectrophotometer (NanoDrop Technologies).

### cDNA synthesis and quantitative RT-PCR

Complementary DNA was synthesized using a reverse transcriptase SuperScript III kit according to the manufacturer’s instructions (Invitrogen). PCR was performed using following primers: Drd2 forward - TACGTGCCCTTCATCGTCAC, Drd2 reverse - CCATTGGGCATGGTCTGGAT, Gapdh forward - ACAGTCCATGCCATCACTGCC, Gapdh reverse - GCCTGCTTCACCACCTTCTTG. qPCR was performed using TaqMan™ Gene Expression Assays (Applied Biosystems) and TaqMan™ probes for Drd2 (Thermo fisher scientific Mm00438545_m1), Adora2a (Thermo Fisher Scientific Mm00802075_m1), Rgs9 (Thermo Fisher Scientific Mm01250425_m1), Gpr52 (Thermo Fisher Scientific Mm03036995_s1), Gpr6 (Thermo Fisher Scientific Mm01701705_s1), Gpr88 (Thermo Fisher Scientific Mm02620353_s1) and Gapdh (Thermo Fisher Scientific Mm99999915_g1). Data was acquired by QuantStudio 3 Real-Time PCR System (Thermo Fisher Scientific). Relativeexpression analysis was performed using data from biological triplicates of each sample byQuantStudio ™ Design and Analysis Software (Thermo Fisher Scientific).

### Affymetrix Mouse Gene 2.0 ST array

Microarray analyses were performed by the CHU de Québec Research Center (CHUL) Gene Expression Platform, Quebec, Canada (www.crchudequebec.ulaval.ca/plateformes/genomic). The quantity of total RNA was measured using an ND-1000 Spectrophotometer (NanoDrop Technologies). Optical density values at 260/280 were consistently above 1.9. The total RNA quality was assayed on an Agilent BioAnalyzer (Agilent Technologies). DNA microarray analyses were carried out with Affymetrix Mouse Gene 2.0 ST arrays according to the manufacturer’s standard protocol.Briefly, total RNA (100 ng per sample) was labeled using the Affymetrix GeneChip^®^ WT Plus Reagent Kit protocol, and hybridized to the arrays as described by the manufacturer (Affymetrix). The cRNA hybridization cocktail was incubated overnight at 45°C while rotating in a hybridization oven. After 16 h of hybridization, the cocktail was removed and arrays were washed and stained in an Affymetrix GeneChip fluidics station 450, according to the Affymetrix-recommended protocol. Arrays were scanned using Affymetrix GCS 3000 7G and Gene-Chip Command Console Software (AGCC) (Affymetrix) to produce probe cell intensity data (CEL). For the quality control, data were analyzed by Affymetrix Expression Console Software to perform background subtraction and normalization of probe set intensities with the method of Robust Multiarray Analysis (RMA). CHP files were imported and analyzed with the Transcriptome Analysis Console Software (Affymetrix).

### Immunohistochemistry

Mice were euthanized 3 weeks after stereotaxic surgery by a lethal dose of ketamine/xylazine and perfused with phosphate buffer saline (PBS) followed by 4% paraformaldehyde (PFA). Brains were incubated in 4% PFA 24h at 4°C. Fixed tissue was sectioned using vibratome (Leica, VT1000S). Next, 40μm sections were washed 3 times for 5 min in PSB. Sections were blocked and incubated with a permeabilization solution containing 5% normal goat serum (Millipore) and 0.5% Triton X-100 (Sigma) in PBS for 2h. Sections were then incubated with primary antibodies diluted in permeabilization solution overnight at 4°C. After three washes in PBS, slices were incubated with secondary antibodies for 2h at room temperature. Sections were rinsed three times for 10 min in PBS before mounting with DAKO mounting medium (DAKO). Staining was visualized using either near-infrared laser scan imaging with an Odyssey Imaging System (LiCor) or confocal microscope (Zeiss LSM 700 or Zeiss LSM 880, Zen 2011 Software).

Following primary antibodies were used: Mouse anti-NeuN (1:500, Millipore MAB377), mouse anti-Calretinin (1:500, Chemicon MAB1578), rabbit anti-Neuropeptide Y (1:500, Sigma N9528), mouse anti-Calbidin (1:500, Sigma C9848), mouse anti-Parvalbumin (1:500, Sigma P3088), rabbit anti-Somatostatin (1:500, Diasoron Inc. 21574), mouse anti-GFP (1:1000, Invitrogen A11120), rabbit anti-Ds Red (1:500, Clontech 632496), mouse anti-HA (1:500, Biolegend MMS-101R-200), rabbit anti-HA (1:500, Sigma H6908), chicken anti-HA (1:500, Abcam Ab9111), rabbit anti-Tyrosine hydroxylase (1:1000, Millipore AB152), rat anti-Dopamine transporter (1:500, Millipore MAB369), anti-norepinephrine transporter (1:1000, Mab technologies NET05-1).

Following secondary antibodies were used: Goat anti-mouse, rabbit, chicken, rat Alexa 488, 568, 647 (1:500, Invitrogen), goat anti-mouse IR Dye 680 (1:10000, Mandel).

### RNAscope in situ hybridization

Fluorescent in situ hybridization (FISH) for Drd2 mRNA (Probe #406501) and Negative Control (Probe #320871) was performed using the RNAscope^®^ Fluorescent Multiplex 2.0 assay as per the manufacturer’s instruction (Advanced Cell Diagnostics, Hayward, CA, USA). Briefly, fresh whole mouse brains were embedded in OCT medium and quickly frozen in isopentane (2-methylbutane) chilled to −80°C. Twenty-micrometer cryosections of brain tissues were then prepared and mounted on SuperFrost Plus slides. Sections were fixed and pre-treated according to the RNAscope^®^ guide for a fresh frozen tissue. After pre-treatment, sections were hybridized with Drd2 and Negative control probes using the HybEZ Hybridization System. After several amplification sets, the sections were counterstained with DAPI and mounted using DAKO. Images were acquired on an LSM880 confocal microscope (Carl Zeiss).

### Statistical analysis

Data are presented as means ± SEM. Two-tailed t test is used in GraphPadPrism 5 software for comparison between two groups (La Jolla, CA) (*p < 0.05, **p < 0.01, ***p < 0.001).

## Results

### The Drd2 positive cell clusters are present throughout the cerebral cortex

RiboTag mice were crossed with BAC D2Cre mice (Gerfen CR *et al.* 2013) resulting in activation of epitope-tagged ribosomal protein Rpl22-HA under the control of its own promoter and its incorporation into ribosomes (Sanz E *et al.* 2009) only in Drd2 expressing cells. This quantitatively dissociates reporter expression level from Drd2 thus providing an all or none-binary signal with greater sensitivity of detections in neurons expressing low levels of receptors.

Immunofluorescent staining against HA on coronal serial sections of D2Cre::RiboTag mice (D2Cre+/RiboHA+ mice, homozygous for floxed Rpl22-HA and heterozygous for D2Cre) revealed Drd2+ cells in previously documented structures such as striatum (STR) (Fig. 1A-D), hippocampus (HIP) (Fig. 1E), thalamus (THAL), hypothalamus (HYP) (Fig. 1C), septum (SPT) (Fig. 1B), VTA and SN (Fig. 1E) supporting the validity of the model. In addition, HA labeled cells were observed throughout the cortex (Fig. 1A-E). Interestingly, several regions such as medial prefrontal cortex (mPFC) (Fig. 1A), claustrum and insular cortex (CL/INS, Fig. 1A-C), cingulate cortex (Cg, Fig. 1B), somatosensory cortex (SS, Fig. 1C), auditory and endopiriform cortices (AUD, EP, Fig. 1D), visual and entorhinal cortices (VIS, Ent, Fig. 1E) contained clusters of reporter-positive cells.

**Figure 1.**
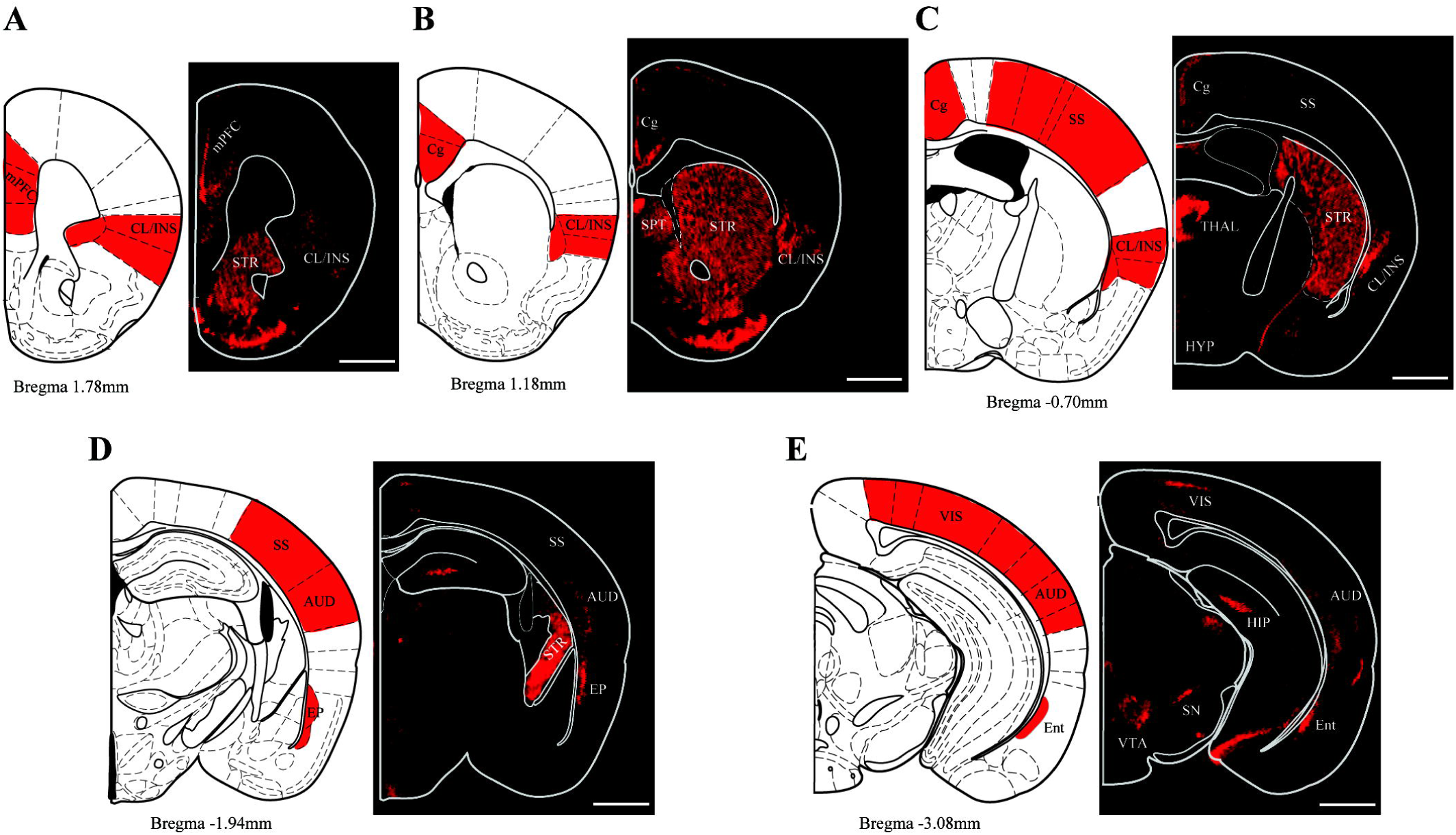
Distribution of Drd2 cell clusters in multiple cortical regions. **(A-E)** Immunofluorescent staining for HA expression reveals neuronal clusters in **(A)** anterior CL/INS and mPFC, **(B)** medial CL/INS and Cg cortex, **(C)** posterior CL/INS and SS cortex, **(D)** EP and AUD cortices, **(E)** Ent and VIS cortices of RiboHA+/D2Cre+ mice. On the left panel, red fields indicate only cortical regions with Drd2+ clusters. Scale bar is 1mm.

D2 EGFP mice are a commonly used BAC transgenic analog reporter system, where GFP is expressed under the control of the Drd2 promoter (Gong S et al. 2003). This links the expression of the reporter to the activity of the promoter, thus limiting GFP detection when the activity of the Drd2 promoter is low. Immunofluorescent staining against GFP was performed on serial coronal sections of D2 EGFP mice to visualize Drd2+ cells in cortical regions and to draw a comparison with the binary RiboTag::D2Cre system. In contrast to RiboTag::D2Cre mice, D2 EGFP mice did not allow for visualization of all identified cortical cluster and showed few cells in mPFC, CL/INS and no cells in, SS, AUD, and VIS (Supplementary Fig. 1). This difference may arise from the high sensitivity of the RiboTag reporter system as compared to D2GFP. However, one caveat of a Cre reporter system is that gene promoters can be transiently activated during brain development (Thompson CL et al. 2014). Thus, it is also possible that Drd2+ cell clusters in different cortical regions of RiboTag::D2Cre mice may consist mostly of cells that were labeled as a result of transient Cre activation during brain development.

### Expression of Cre recombinase in Drd2 cells during adulthood

To rule out the possibility of transient embryonic Cre expression resulting in false positive HA-labeling, Cre expression needs to be demonstrated during adulthood. Detection of Cre by antibodies can be difficult if the recombinase is expressed at low levels in some Drd2 cell. Thus, a Cre dependent DIO-mCherry reporter virus was used to confirm Cre activity and to establish that HA labeling did not result from Cre expression during brain development. A second Cre independent EGFP virus was used as a control for viral infection. AAV hSyn-DIO-mCherry and AAV hSyn-EGFP viruses were mixed in a 1:1 titer ratio and injected into cortical areas containing clusters of HA-positive cells in D2Cre+RiboHA+ mice. Because of the complex anatomy of CL/INS, this structure was arbitrarily divided into anterior (CL/INS ant, level of Fig. 1A), medial (CL/INS med, level of Fig. 1B) and posterior (CL/INS post, level of Fig. 1C) regions. GFP, mCherry, and HA were visualized by immunostaining and confocal microscopy. HA+ staining revealed all the cells that have or had Cre activity, mCherry+ cell labeling corresponded to cells where Cre is active in adulthood while GFP+ cells labeling represented all the cells that were infected with viruses. This triple labeling allowed for the quantification of the co-infection rate of mCherry+ cells by GFP expressing virus (mCherry+GFP+ out of all mCherry+) and the percentage of HA+ cells expressing Cre in adulthood (mCherry+HA+GFP+ out of all HA+GFP+) (Fig. 2A). Co-infection rate of mCherry+ cells by GFP expressing virus was close to 80% in most cortical regions (Fig. 2B). This supports the use of GFP expression as a reliable marker for virus-infected cells. Furthermore, 80 to 100% of virus-infected HA+ cells (HA+/GFP+) were also mCherry+ (mCherry+/HA+/GFP+) depending of cortical region (Fig. 2B). Overall, this indicates that more than 80% of reporter-labeled (HA+) cells also display Cre recombinase activity during adulthood, thus confirming that Rpl22-HA expression is representative of adult Cre expression in these cortical areas.

**Figure 2.**
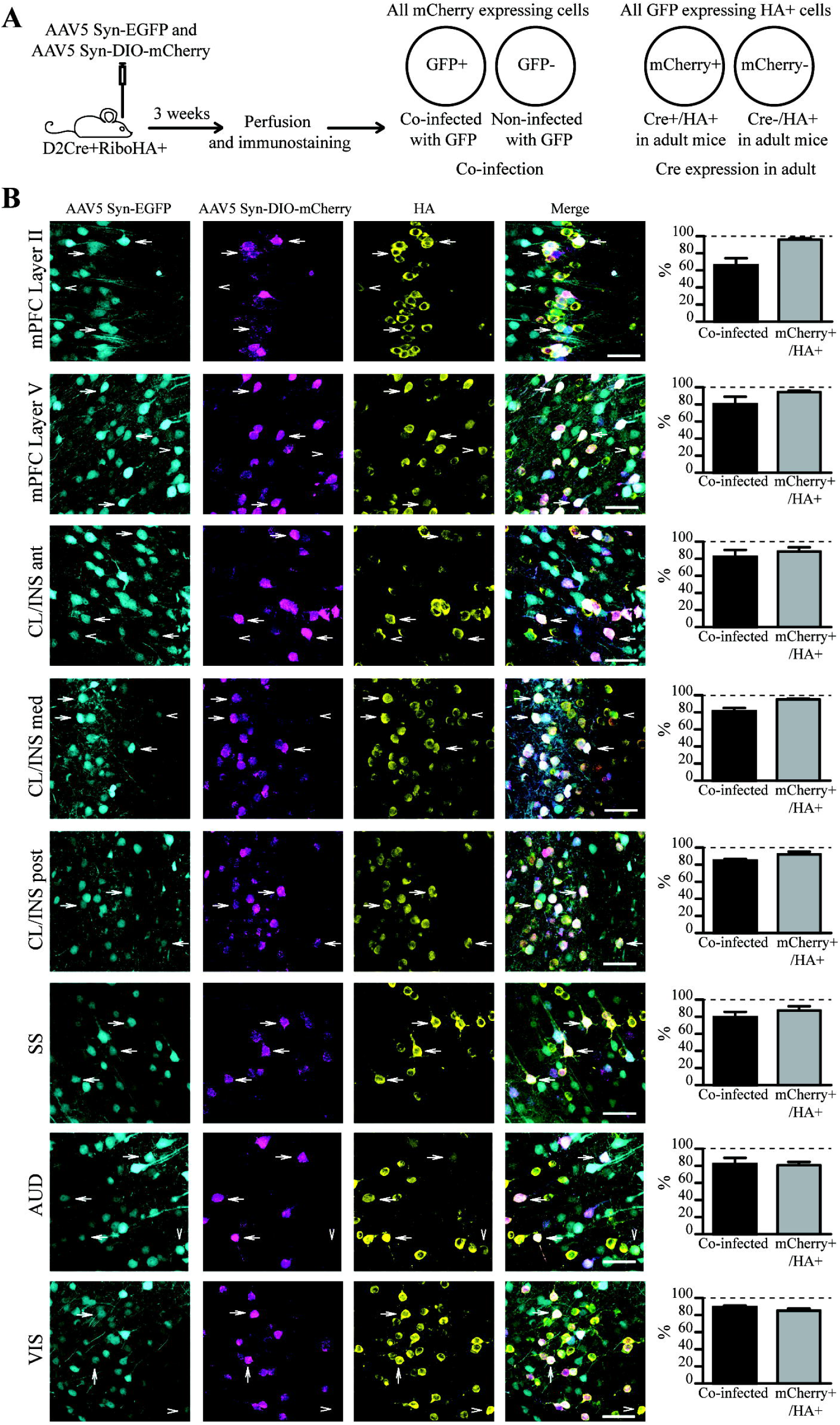
Cortical clusters of HA+ cells express Cre during adulthood. **(A)** Schematic representation of the experimental design. **(B)** Immunostaining of HA in brain slices of RiboHA+/D2Cre+ mice injected with the mixture of AAV Syn-EGFP and AAV Syn-DIO-mCherry viruses. Right panel shows quantification of the % of co-infection (i.e. % of EGFP+/ mCherry+ cells out of all mCherry expressing cells) and % of Cre expression in HA+ cells (i.e. % of mCherry+/EGFP+/HA+ cells out of all EGFP+/HA+ cells) (n=3 mice per brain region). Arrows show mCherry+/EGFP+/HA+ cells, arrowheads show mCherry-/EGFP+/HA+ cells. Scale bar is 50μm.

### Drd2 mRNA is expressed in multiple cortical regions

To demonstrate and quantify Drd2 expression directly in reporter-labeled cells (HA+), HA-tagged ribosomes were immunoprecipitated (RiboTag-IP) from the different HA positive cortical clusters. For qualitative PCR, primers that amplify from exon 4 to 6 of the Drd2 transcript were designed, allowing to distinguish between D2S (missing exon 5) and D2L isoforms. RiboTag-IP from the STR of D2Cre+ RiboHA+ and D2Cre-RiboHA+ mice followed by RT-PCR was carried out to validate the specificity. A major band corresponding to D2L and a minor band corresponding to D2S isoforms were detected in all conditions except in RiboTag-IP from Cre negative mouse, where Drd2 and housekeeping gene Gapdh were absent (Fig. 3A). RiboTag IP followed by RT-PCR was performed for all cortical clusters. Samples obtained from STR and HIP were included as positive controls while liver-derived samples were employed as a negative control. D2L was the predominant isoform detected by PCR, although D2S was also expressed to a lesser extent in all Drd2+ brain regions, while expression of both isoforms was undetectable in liver (Fig. 3B).

**Figure 3.**
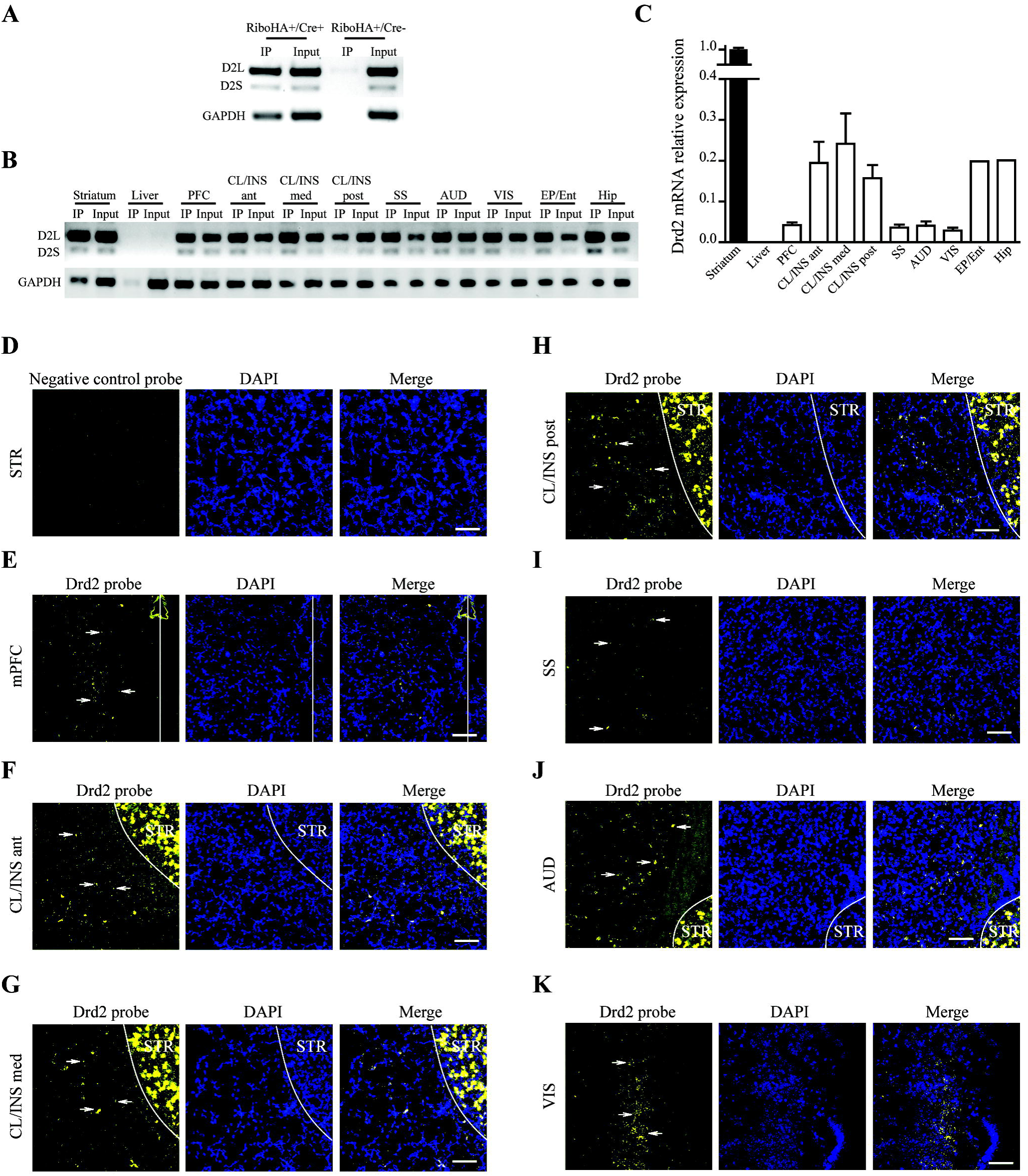
Expression of Drd2 mRNA in multiple cortical regions. **(A)** Immunoprecipitation of polysomes followed by RT-PCR for Drd2 from striatum of RiboHA+/Cre+ and RiboHA+/Cre-mice (pool of 10 mice per genotype). **(B-C)** Immunoprecipitation of polysomes followed by **(B)** RT-PCR (pool of 10 mice per brain region) or **(C)** RT-qPCR for Drd2 from multiple brain regions and liver (n=4 samples per brain region, each sample consists of RNA pooled from 5 mice). **(D-K)** RNAscope^®^ *in-situ* hybridization for Drd2 and a negative control probe on mouse brain slices. Note that every little point represents a mRNA of Drd2, highlighted by the arrow. Scale bar is 100μm.

For quantitative PCR (RT-qPCR), validated TaqMan™ probes were used to quantify the recruitment of Drd2 (D2L and D2S) mRNA to ribosomes specifically in Drd2+ cells from the different brain regions (Fig. 3C). In most cortical areas, RiboTag positive cells displayed 5-10% of ribosome recruited Drd2 mRNA as compared to Drd2 expressing cells from the STR. In CL/INS, EP/Ent and HIP levels of ribosome-associated Drd2 mRNA reached ~20-25% (Fig.3C).

The presence of Drd2 mRNA was further validated by using enhanced fluorescent *in situ* hybridization for RNA (RNAscope^®^) on coronal sections of the adult mouse brain. This technology allows very high amplification of the signal with virtually no background, thus making it possible to visualize mRNAs down to a single-molecule level (Wang F et al. 2012; Tubbs RR et al. 2013). Expression of Drd2 mRNA was detected in all the identified cortical clusters, while negative control probe showed no signal, even in the STR (Fig. 3 D-K). Taken together with results obtained from immunoprecipitated ribosomes, this indicates that Drd2 mRNAs are present in all HA positive cortical regions identified in the adult mouse brain.

### Molecular heterogeneity of Drd2 cells across different cortical regions

Striatal Drd2 cells have been shown to co-express different genes that can sometimes be used as proxies for Drd2 expression. To compare the gene expression profiles between Drd2 cells of STR and various cortical regions several transcripts were selected. GPCRs, such as Adora2A, Gpr88, Gpr52, and Gpr6 have been shown to be the most enriched in the striatum, with Gpr88 expressed in Drd1 and Drd2 neurons, while Gpr52, Gpr6, and Adora2A are specifically expressed in Drd2, but not Drd1 striatal neurons (Komatsu H et al. 2014). The regulator of G protein signaling 9 (Rgs9), is a member of the RGS family of GTPase accelerating proteins. RGS9 is known to be expressed in Drd2 neurons of the striatum, where it participates in the regulation of Drd2 G-protein-mediated signaling (Rahman Z et al. 2003; Kovoor A et al. 2005). RNA was immunoprecipitated from various cortical regions of D2Cre+RiboHA+ mice. RT-qPCR on immunoprecipitated (RiboTag IP) RNA and Total (input) RNA was performed using TaqMan™ probes for Drd2, Adora2A, Gpr88, Gpr52, Gpr6 and Rgs9 (Fig. 4A). The relative expression of the Drd2 transcript in RiboTag IP RNA was significantly higher compared to Total RNA in all regions. This confirms that the Drd2 transcript is enriched in RiboTag IP samples (Fig. 4B). However, the picture was different for other targets across various regions. Adora2A transcript was enriched in Drd2 cells of CL/INS, SS and STR (Fig. 4C). Gpr88 was enriched in Drd2 cells of CL/INS and STR (Fig. 4D). The Gpr52 transcript was enriched in Drd2 cells of CL/INS, SS and STR (Fig. 4E). The Gpr6 transcript was enriched in Drd2 cells of mPFC, CL/INS, SS, AUD and STR (Fig. 4F). The Rgs9 transcript was enriched in Drd2 cells of mPFC, CL/INS, SS, AUD, VIS but not STR (Fig. 4G). Overall this shows a level of molecular heterogeneity of Drd2 cells across various cortical regions.

**Figure 4.**
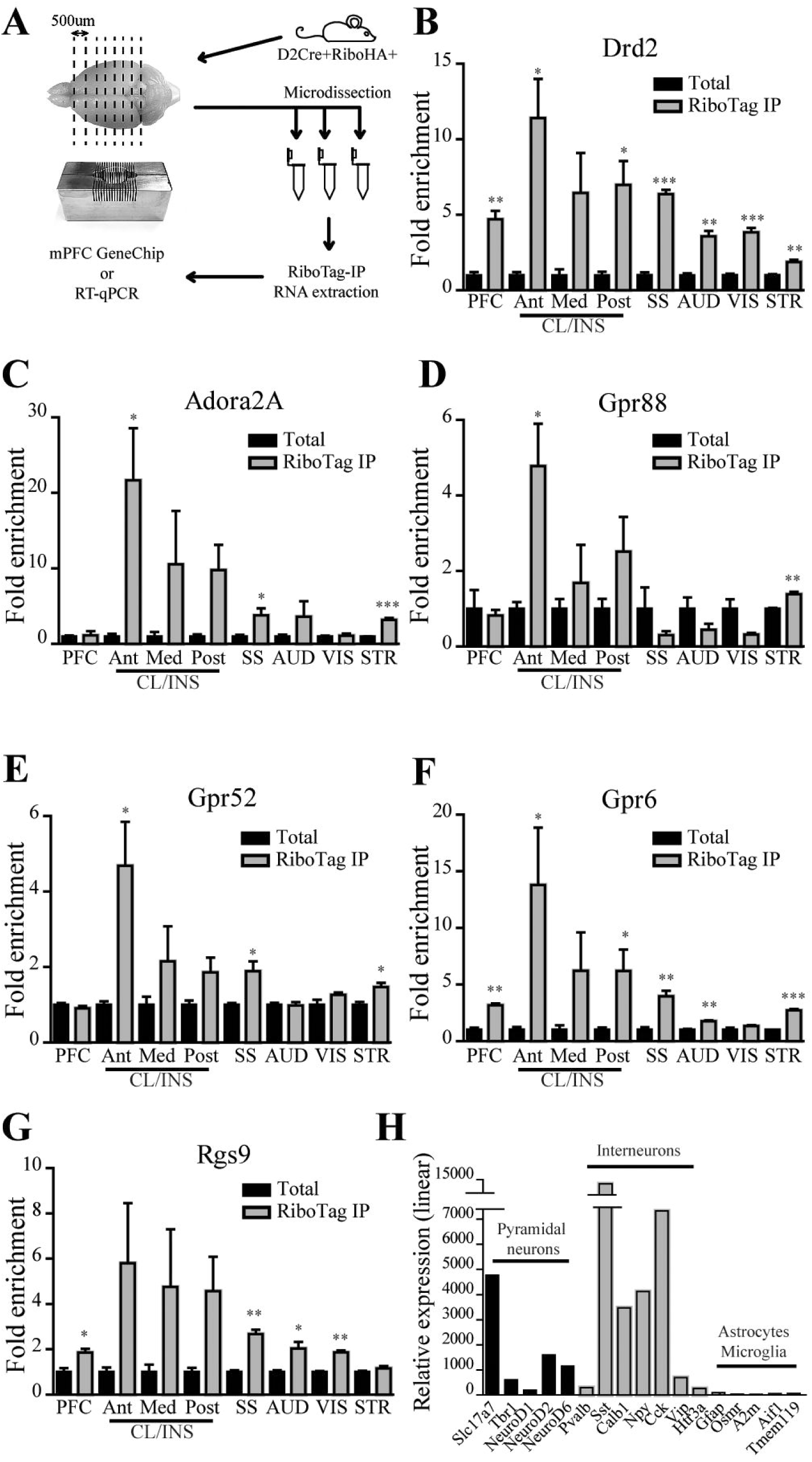
Transcriptomic heterogeneity of Drd2 cells from different cortical regions. **(A)** Schematic representation of the experimental design. **(B-G)** RT-qPCR of mRNA isolated by HA immunoprecipitation from multiple cortical regions of D2Cre::RiboTag mice. Fold enrichment of RiboTag IP RNA and total RNA for **(B)** Drd2, **(C)** Adora2A, **(D)** Gpr88, **(E)** Gpr52, **(F)** Gpr6, **(G)** Rgs9 (n=3 samples per brain region, each sample consists of RNA pooled from 5 mice). **(H)** Relative expression of various transcripts from GeneChip analysis of RiboTag IP RNA from mPFC Drd2 cells (pool of 5 mice).

A more complete characterization of a translatome profile of Drd2+ cells of mPFC was performed considering the high interest given to Drd2+ cells of this region. The ribosomes were immunoprecipitated from dissected cortices of D2Cre+RiboHA+ mice (Fig. 4A). Gene expression analysis was carried out using Affymetrix mouse GeneChip2.0 ST to reveal the complete translatome of mPFC Drd2+ cells (Supplementary Table 1, GSE: 115887). These cells show high expression of pyramidal neuron markers: Vesicular glutamate transporter 1 (Vglut1 or Slc17a7), T-box brain gene 1 (Tbr1), neurogenic differentiation factor 1, 2, and 6 (NeuroD1, NeuroD2, NeuroD6) (Hevner RF et al. 2006; Molyneaux BJ et al. 2007). Interneuron transcripts were also highly expressed: somatostatin (Sst), parvalbumin (Pvalb), Calbindin (Calb1), Neuropeptide Y (Npy), cholecystokinin (Cck), vasoactive intestinal polypeptide (Vip), 5-hydroxytryptamine (serotonin) receptor 3A (Htr3A) (Rudy B et al. 2011; Taniguchi H 2014; Yavorska I and M Wehr 2016). In contrast, astrocyte (Gfap, Osmr, A2m) (Ito K et al. 2016) or microglia (Aif1 or Iba1, Tmem119) (Bennett ML et al. 2016) markers showed very low expression (Fig. 4H, Supplementary Table 1). This indicates that mPFC Drd2 cells represent a heterogeneous pool of neurons and interneurons with virtually no astrocytes and microglia.

### Cellular heterogeneity of Drd2 clusters in different cortical regions

Translational profiling data suggests that Drd2 is expressed in principal neurons and in several interneuron subtypes of mPFC. Double immunofluorescent staining of HA in combination with several cell type selective markers was thus performed in brain slices of D2Cre+RiboHA+ mice to identify cell types that express Drd2 across all the cortical clusters and quantify their representation in the total pool of the cells in each region. NeuN and HA co-labeling confirmed that all Drd2 expressing cells are neurons (Fig. 5A, B).

**Figure 5.**
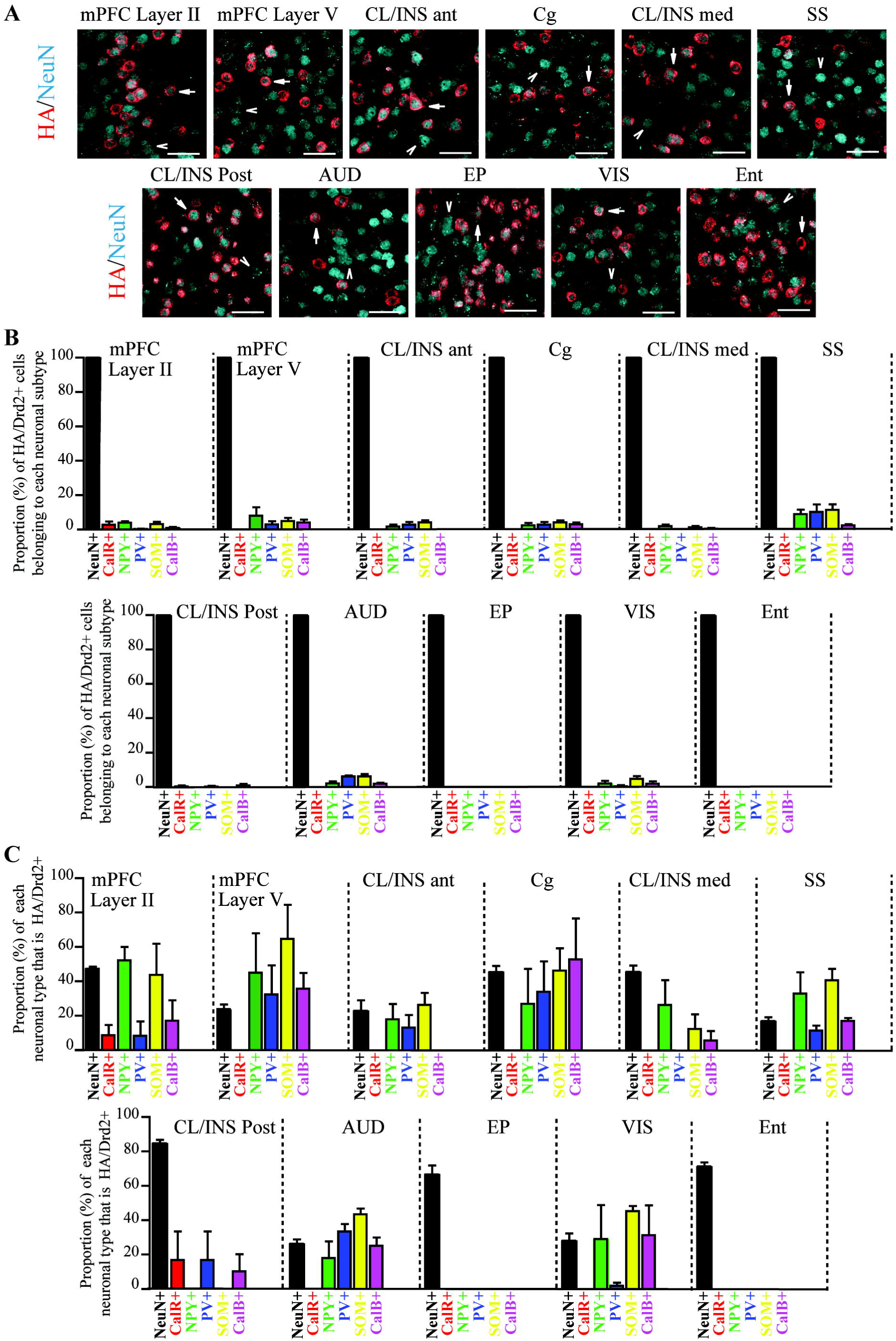
Drd2 expressing cells throughout the cortex. **(A)** Immunofluorescent staining for HA and NeuN in different cortical regions of RiboHA+/D2Cre+ mice. **(B)** Quantification of proportion (%) of HA+ cells belonging to each neuronal subtype. **(C)** Quantification of proportion (%) of each neuronal subtype that is HA+. n=3 mice per brain region. Scale bar is 40μm.

Five different phenotypic markers were then used to identify interneuron subtypes that express Drd2: calretinin (CalR), neuropeptide Y (NPY), PV, somatostatin (SOM) and calbindin (CalB) (Fig. 5B, C, and Supplementary Fig. 2). Interneurons accounted only for a small portion of Drd2+ cells in all cortical regions (Fig. 5B, Supplementary Fig. 2). This suggests that the vast majority of Drd2 expressing cells are principal neurons.

Assuming that high representation of Drd2+ cells could be indicative of an important functional implication, the proportion in each cell subtype was quantified. Depending on the region, 20-85% of all NeuN positive cells expressed Drd2 (Fig. 5C). Drd2+ neurons were represented unevenly in various interneuron subtypes across cortical regions (Fig. 5C, Supplementary Fig. 2). For example, 30-50% of different types of interneurons in mPFC layer II, V, Cg, SS, AUD, and VIS also expressed Drd2, few interneurons expressed Drd2 in CL/INS while in regions like EP and Ent, almost no Drd2+ interneurons were detected (Fig. 5C, Supplementary Fig. 2). Overall, this indicates that Drd2 is expressed in different heterogeneous neuronal populations and that these populations vary across cortical regions.

### The dopaminergic input of Drd2 cells from different cortical regions

We then investigated whether cortical clusters of Drd2 cells receive dopaminergic input. Brain sections were immunostained for tyrosine hydroxylase (TH) and dopamine active transporter (DAT) to detect dopamine neuron terminals. TH labeled fibers were detected in all regions, with the highest density and labeling intensity in CL/INS and STR (Fig. 6A). In contrast, DAT labeled fibers were only visible in STR (Fig. 6B). The absence of DAT staining in cortical TH positive termini has previously been reported (Sesack SR 2014), suggesting that in absence of DAT other monoamine transporters can participate in dopamine reuptake (Morón JA et al. 2002; Smith CC and RW Greene 2012). Indeed, we have detected the presence of norepinephrine transporter (NET) in all the cortical regions. Moreover, a fraction of TH+ fibers in those regions were also NET+ (Supplementary Fig. 3).

**Figure 6.**
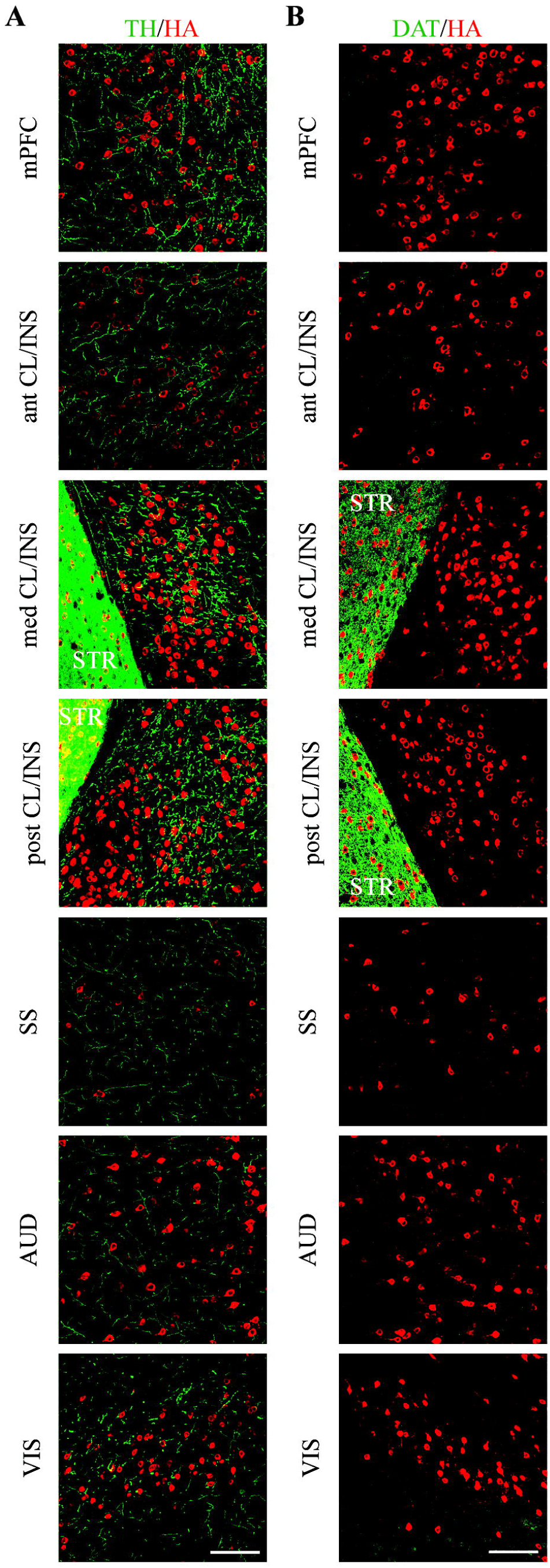
Dopaminergic projections in cortical Drd2 clusters. Immunofluorescent staining for HA (red, Drd2 cells) and **(A)** TH (green, axons), **(B)** DAT (green, axons) in RiboHA+/D2Cre+ mice. Scale bar is 100μm.

### Projection targets of Drd2 cells from different cortical regions

The direct projection targets of Drd2+ cortical neurons were mapped to characterize the position of Drd2+ cell clusters in the wider context of brain connectivity. AAV hSyn-EGFP and AAV hSyn-DIO-mCherry viruses were mixed and injected into different cortical regions in D2Cre mice (Fig. 7, Supplementary Fig. 4-9). GFP and mCherry were visualized on coronal serial sections by confocal microscopy to identify: Drd2+ neurons that are labeled by both GFP and mCherry and appear yellow, and Drd2 negative neurons (Drd2-) that are labeled only by GFP (Fig. 7, Supplementary Fig. 4-9). Maps of Drd2+ and Drd2-neuron projections from cortical clusters were created (Fig. 7, Supplementary Fig. 4-9). High magnification images resolving cell bodies or axonal varicosities were taken to verify the injection and projection sites respectively (Fig. 7, Supplementary Fig. 4-9). Projections from all regions are schematically summarized in Fig. 8.

**Figure 7.**
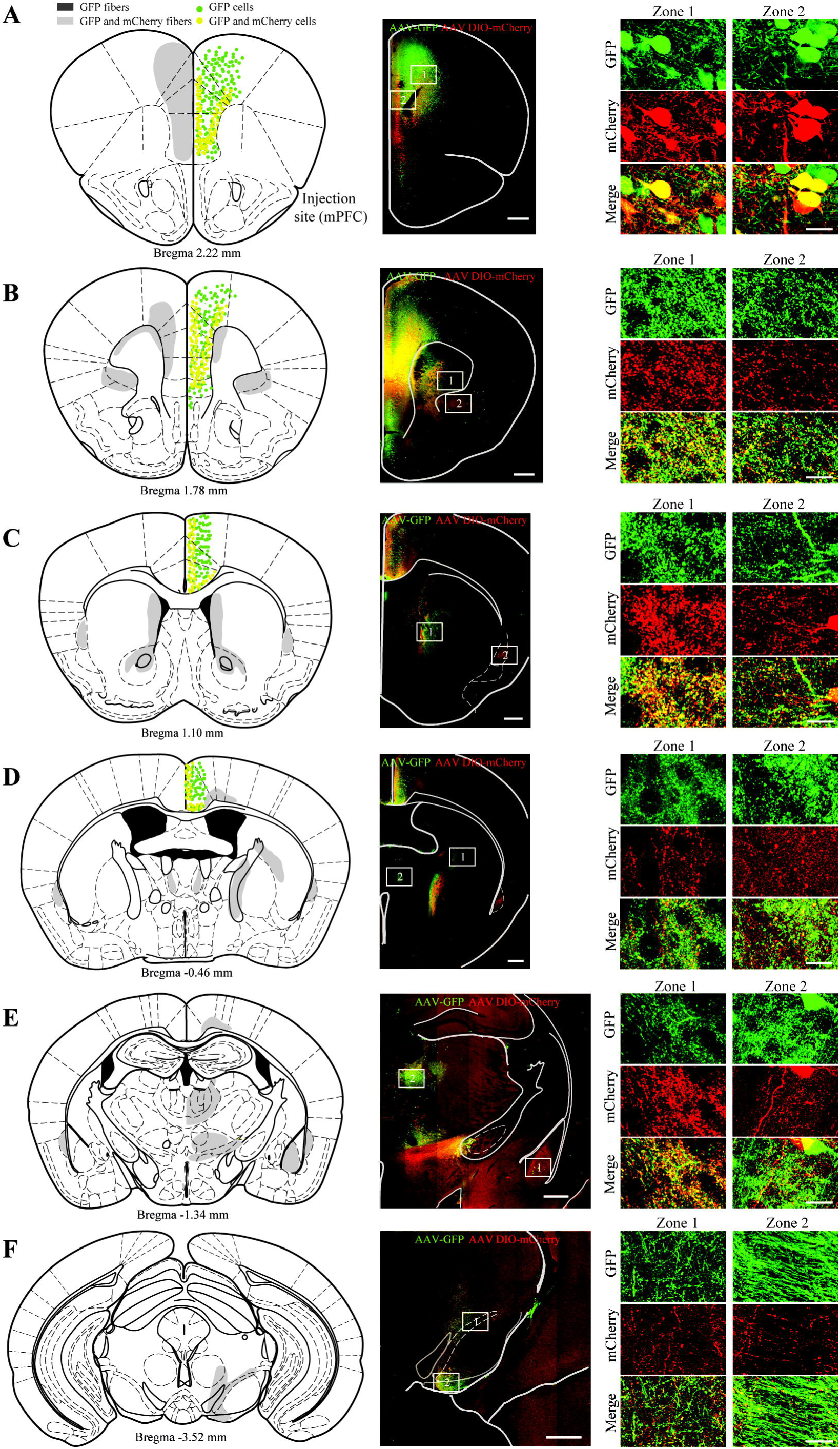
Projections of mPFC Drd2 neurons. **(A-F)** Immunofluorescent staining for GFP and mCherry (middle and right panel) and graphical representation of injection and projection sites (left panel). Colored dots represent injection sites and virus-infected cell bodies (green: infected with AAV GFP, yellow: infected with AAV GFP and AAV DIO-mCherry). Colored fields represent projection areas of infected neurons (dark grey: projections of only AAV GFP infected cells, light grey: mixed projections of AAV GFP/AAV DIO-mCherry and AAV GFP infected cells), scale bar is 500μm. Right panel Zone 1 and Zone 2 represent high magnification images of corresponding boxes of the middle panel, scale bar is 20μm.

**Figure 8.**
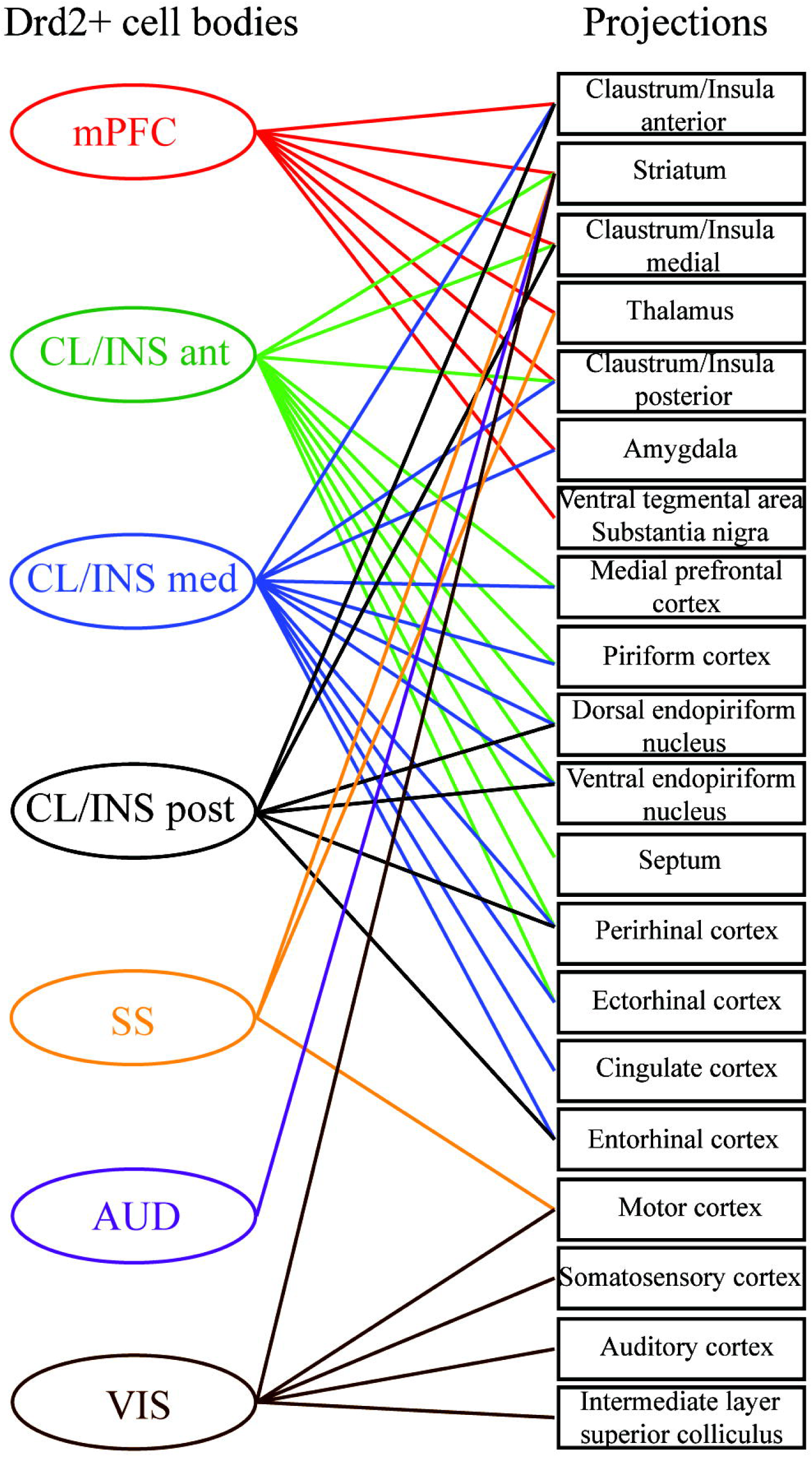
Schematic representation of projections of Drd2 expressing cortical neurons. Drd2+ cell projections from different cortical regions (shown in Fig.7 and Supplementary Fig. 4-9) are summarized.

For the mPFC, Drd2+ and Drd2-neurons project to the same targets: contralateral mPFC (Fig. 7A-B), CL/INS (Fig. 7B-D), STR (including nucleus accumbens) (Fig. 7B-D), THAL (Fig. 7D-E), amygdala (AMY) (Fig. 7E) and SN/VTA (Fig. 7F). In contrast, other regions showed only partial overlaps.

For the CL/INS ant, Drd2+ and Drd2-neurons project to mPFC (Supplementary Fig. 4A-B), CL/INS (Supplementary Fig. 4B-E), STR (Supplementary Fig. 4C), piriform cortex (Pir), dorsal (DEn) and ventral (VEn) endopiriform nucleus (Supplementary Fig. 4B-E), SPT (Supplementary Fig. 4C), ectorhinal (Ect) and perirhinal (PRh) cortices (Supplementary Fig. 4F). However, in contrast to Drd2-, Drd2+ neurons do not project to controlateral mPFC and CL/INS ant (Supplementary Fig. 4A), controlateral CL/INS post (Supplementary Fig. 4D), controlateral VEn, DEn, Ect, PRh (Supplementary Fig. 4E-F), certain parts of THAL and HYP (Supplementary Fig. 4E) and SN/VTA (Supplementary Fig. 4F).

For the CL/INS med, Drd2+ and Drd2-neurons project to mPFC, CL/INS, Pir, DEn, VEn, (Supplementary Fig. 5A-D), Ent, PRh, Ect (Supplementary Fig. 5E-F), Cg (Supplementary Fig. 5C-D) and AMY (Supplementary Fig. 5E). However, Drd2+ neurons do not project to controlateral mPFC, INS (Supplementary Fig. 5A) and SN (Supplementary Fig. 5F).

For the CL/INS post, Drd2+ and Drd2-neurons project to CL/INS (Supplementary Fig.6A-D), DEn, VEn, PRh, Ent (Supplementary Fig. 6D, F). However, Drd2+ neurons do not project to mPFC (Supplementary Fig. 6A-B) and STR (Supplementary Fig. 6C, E).

For the SS, Drd2+, and Drd2-neurons project to motor (M) cortex (Supplementary Fig. 7A-D), STR (Supplementary Fig. 7D-E) and certain thalamic nuclei (Supplementary Fig. 7E). However, Drd2+ neurons do not project to certain regions of THAL (Supplementary Fig. 7E-F).

For the AUD, Drd2+, and Drd2-neurons project to STR (Supplementary Fig. 8E).However, Drd2+ neurons do not project to THAL (Supplementary Fig. 8E-F).

For the VIS, Drd2+, and Drd2-neurons project to STR (Supplementary Fig. 9B-D), M, SS, AUD (Supplementary Fig. 9B-D, F), an intermediate gray layer of the superior colliculus (InG) (Supplementary Fig. 9F). However, Drd2+ neurons do not project to certain thalamic nuclei (Supplementary Fig. 9D-E).

## Discussion

We used a highly sensitive, multimodal approach to map Drd2 expressing neurons in a cortex-wide fashion. The use of a binary reporter mouse and a Cre dependent virus uncovered previously uncharacterized clusters of Drd2+ cells in mPFC, CL, SS, AUD, and VIS. The presence of Drd2 mRNA was qualitatively and quantitatively demonstrated across all cortical clusters. Transcript enrichment analysis, as well as translational profiling and immunolabeling with cell-specific markers, revealed molecular and cellular heterogeneity of Drd2+ cells across the cortex. Finally, we constructed a map of Drd2 expressing neuron projections for all cortical clusters, providing the general framework of connectivity to guide functional investigations.

Along with cortical clusters, Drd2 expression was detected also in previously characterized structures such as in STR, SN, VTA, HYP, SPT, superior colliculus, THAL, HIP and AMY (Fig. 1) (Missale C *et al.* 1998; Gerfen CR 2000; Vallone D et al. 2000; Rieck RW et al. 2004; Seeman P 2006; Bolton AD et al. 2015; Li Y et al. 2015; Puighermanal E *et al.* 2015; De Bundel D et al. 2016). The presence of Cre recombinase in reporter-labeled cells of adult mouse cortex confirmed Drd2 promoter activity during adulthood (Fig. 2). For all cortical regions, Drd2 mRNA expression was confirmed quantitatively by RT-qPCR on ribosome-associated mRNAs as well as qualitatively by fluorescent *in situ* hybridization on adult mouse brain slices (Fig. 3). All these results point to the specificity, sensitivity, and selectivity of this model.

Along with Drd2, we selected Gpr52, Gpr88, Gpr6, Adora2a and Rgs9 transcripts to study their enrichment in ribosome bound RNA from Drd2+ cells versus total mRNA from each region. GPCRs Gpr52, Gpr88, Gpr6, and Adora2a are the most enriched GPCR transcripts in STR (Komatsu H *et al.* 2014) and are considered potential therapeutic targets for neuropsychiatric disorders (Komatsu H 2015). Rgs family protein 9 is a regulator of G protein signaling in part by stimulating the GTPase activity of the G protein α subunits. Rgs9 has been shown to localize to striatal Drd2+ neurons and regulates Drd2 receptors signaling (Rahman Z *et al.* 2003; Celver J et al. 2010). As expected, all the regions showed enrichment for the Drd2 transcript (Fig. 4). In line with previous studies, STR Drd2+ cells were enriched for Adora2A, Gpr88, Gpr52 and Gpr6 (Komatsu H *et al.* 2014) (Ferré S et al. 2008). STR Drd2+ cells did not show enrichment for Rgs9, indicating that presence of Rgs9 in the STR is not exclusive to Drd2+ neurons. In contrast, Drd2+ cells of cortical clusters show enrichment in Rgs9 indicating its possible selective functional involvement in Gia/o mediated Drd2 signaling in those neurons (Fig. 4).

Apart from STR, Adora2A was enriched in Drd2 positive neurons in CL/INS, SS, and AUD, but not mPFC and VIS. Adora2A is known to be co-expressed with Drd2 in STR and Adora2A Cre mice are often used as a proxy for Drd2+ cells (Schiffmann SN et al. 1991; Schiffmann SN and JJ Vanderhaeghen 1993). However, our results show that it is not the case for mPFC and VIS, where Adora2A is not enriched, thus not exclusively co-expressed, in Drd2+ neurons. Gpr88 was enriched in CL/INS and Gpr52 in CL/INS and SS Drd2+ neurons. In addition to CL/INS and SS, Gpr6 was also enriched in mPFC and AUD Drd2+ cells. Overall, cortical Drd2+ cells are heterogeneous with regard to expression of studied striatum enriched transcripts. This indicates that therapeutic and genetic targeting of those GPCRs may not be specific for striatum or striatal Drd2+ neurons and can impact various cell types throughout the cortex. This underscores not only possible limitations of therapeutic interventions targeting these receptors, but also suggests strategies to modulate the activity of Drd2+ cells subpopulations more specifically.

Along with molecular heterogeneity, Drd2+ cells are also heterogeneous in regard to cell types. GeneChip analysis of ribosome-bound mRNAs established a translational profile of mPFC Drd2+ neurons (Fig. 4, Supplementary Table 1). mPFC Drd2+ neurons express markers for pyramidal cells (Vglut1, Tbr1, NeuroD1, NeuroD2, NeuroD6) (Hevner RF *et al.* 2006; Molyneaux BJ *et al.* 2007) and 3 major independent subtypes of cortical interneurons (PV, SOM, Htr3A) (Rudy B *et al.* 2011; Taniguchi H 2014; Yavorska I and M Wehr 2016), but not for glial cells, once again indicating their cellular heterogeneity (Supplementary Table 1).

Previous studies have suggested that in the mPFC, Drd2 is expressed in a small population of pyramidal neurons (5-25%) and interneurons (5-17%) with most of the Drd2+ interneurons being PV+ (Le Moine C and P Gaspar 1998; Santana N *et al.* 2009; Fitzgerald ML et al. 2012; Tritsch NX and BL Sabatini 2012). However, our characterization of Drd2 expressing cell types by immunofluorescent staining with multiple markers in mPFC and other cortical regions revealed that in mPFC 50% of layer II and 30% of layer V pyramidal neurons express Drd2 (Fig. 5, Supplementary Fig. 2). Moreover, up to 40% of interneurons (depending on subtype) express Drd2 (Fig. 5, Supplementary Fig. 2). Interestingly, expression of Drd2 was not limited to PV+ cells but was more frequently observed in other subtypes of interneurons, mostly NPY+ and SOM+ (Fig. 5, Supplementary Fig. 2). This data is in line with our translatome analysis (Fig. 4H, Supplementary Table 1).

Depending on brain regions, a significant portion of SOM+ and PV+ interneurons may also express CalR, CalB and NPY markers. However, it has been shown that only a small fraction of PV+ cells express SOM marker, thus PV+ and SOM+ interneurons represent two independent groups of interneuron with minimal overlap (Rudy B *et al.* 2011; Taniguchi H 2014; Yavorska I and M Wehr 2016). Furthermore, Drd2+ cells were distributed unevenly in different Drd2+ cortical clusters with a proportion among NeuN+ cells ranging from 20% to 85% (Fig. 5, Supplementary Fig. 2). The proportion of Drd2+ cells among different types of interneurons also varied among cortical brain regions (Fig. 5, Supplementary Fig. 2). This indicates that PV+ cells cannot account for a majority of Drd2+ interneurons in the mPFC or other cortical regions.

By using AAV mediated tracing, we constructed a general map of the Drd2+ neuron projections (Fig. 7, 8, Supplementary Fig. 4-9). Our results show that Drd2+ neurons do not always follow the characteristic pattern of projection of the region where they reside. These results may point to the possible function of Drd2+ cells of a certain brain region. However, more detailed tracing for each region has to be performed. For instance, by quantitatively comparing the innervations of a given region by Drd2+ cells to the innervations by other cells from the same brain area. This may indicate whether an area is preferentially innervated by Drd2+ cells, thus providing deeper insight into the role of Drd2+ cells in a given network.

When analyzing the presence of dopaminergic fibers, we noted that all the regions receive TH immunoreactive innervation, while DAT staining was undetectable (Fig. 6). All the cortical regions were also innervated to various degrees by NET+ fibers. Moreover, a fraction of TH+ fibers were also NET+ (Supplementary Fig. 3). This can be indicative for: a) TH positive fibers are dopaminergic fibers from VTA or SN that do not express DAT (Morón JA *et al.* 2002; Smith CC and RW Greene 2012; Sesack SR 2014). b) TH positive fibers can also be NET+ noradrenergic fibers from locus coeruleus that can also co-release dopamine (Devoto P et al. 2005, 2005; Smith CC and RW Greene 2012; Kempadoo KA et al. 2016). c) Drd2+ cells do not receive dopaminergic innervations on their cell bodies. Some of Drd2 cell clusters, such as mPFC, SS, AUD and VIS project to STR, and it is thus possible that Drd2 in these neurons could be located in axonal terminals in STR, where they may sense dopamine (Wang H and VM Pickel 2002; Pinto A and SR Sesack 2008; Sesack SR 2014).

Our investigation revealed the existence of clusters of Drd2+ neurons in the CL/INS, SS, AUD, and VIS (Fig. 1). Interestingly, certain symptoms of schizophrenia can be linked to these cortical regions. Multiple forms of hallucinations; auditory verbal (van der Gaag M 2006; Javitt DC and RA Sweet 2015), visual and multisensory (David CN et al. 2011) indicate a possible involvement of SS, AUD and VIS cortices. Delusions in schizophrenia have also been linked to CL/INS (Cascella NG et al. 2011; Patru MC and DH Reser 2015). Hallucinations and delusions are a part of positive symptoms of schizophrenia. Drd2 is a primary pharmacological target for the treatment of positive symptoms (Rolland B et al. 2014) as well as delusion therapies in schizophrenic and non-schizophrenic (bipolar disorder, depression) psychiatric disorders (Patru MC and DH Reser 2015). Furthermore, the molecular and functional heterogenicity of Drd2+ neurons should be taken into consideration and can provide opportunities for therapeutic strategies targeting theses neuronal populations. Overall, our results thus pave the way for a thorough functional reexamination of cortical Drd2 in rodents and humans, which could provide valuable information about neuronal circuits involved in psychotic and mood disorders.

## Acknowledgments

JMB is Canada Research Chair in Molecular Psychiatry. This work was supported by a grant from Canada Institutes of Health Research (CIHR, MOP-136916) to JMB. JMB is NARSAD independent investigator and One-Mind Rising Star awardee.

